# Single-cell RNA-seq reveals conserved and divergent cellular states across wound types and species

**DOI:** 10.64898/2026.07.07.737032

**Authors:** Dahim Choi, Mojtaba Bakhtiari, Aastha R. Amin, Joshua Mann, Swati Bhasin, Manoj K. Bhasin

**Author notes:** ***Corresponding author:*** Manoj K. Bhasin, MS, PhD, Emory University, Department of Pediatrics HSRB-II, Room S335, 1760 Haygood Drive, Atlanta, GA, 30322.

## Abstract

Chronic wounds, such as diabetic foot ulcers, fail to progress through the normal healing process and impose a significant burden on healthcare systems. While previous single-cell studies have characterized specific wound conditions, a unified understanding of the shared and distinct cellular landscapes across diverse wound microenvironments has been lacking. Therefore, we integrated over 500,541 cells from patients and mice across multiple wound conditions, including acute wound, diabetic foot ulcer, and venous ulcer as well as their healing outcome. Fibroblast-focused analysis identified a bifurcation in differentiation trajectories and identified STAT3 as a potential regulator of a reparative program in chronic wound. Furthermore, we discovered immune dysfunctions in non-healed chronic wounds, contrasting the quiescent memory-like T cells and TIMP1+ macrophages in healed chronic wounds with the exhausted T cells and foamy SPP1+ macrophage enriched in non-healed chronic wounds. Finally, we translated these results into a clinically applicable three-gene signature (CHI3L1, TIMP1, and SPP1) that accurately predicts chronic wound healing. To support wound biology community, we developed WoundSCAtlas, an interactive web resource for exploring diverse wound pathologies. In conclusion, this work provides a comprehensive and cross-species landscape of chronic wound healing, identifying conversed wound outcome-associated molecular programs, predictive biomarker, and interactive data resource.

## Introduction

Chronic wounds, including diabetic foot ulcers (DFU) and chronic venous ulcers (CVU), remain as a serious clinical challenge worldwide (McDermott et al., 2023, Probst et al., 2023). Unlike acute wounds, which typically go through progressive wound healing stages including hemostasis, inflammation, proliferation, and remodeling, the chronic wounds do not show such progression. As a result, they not only impose an economic burden on patients as well as healthcare systems but also impair patients’ quality of life (Armstrong et al., 2020, Olsson et al., 2019, Pastar et al., 2023, Zhao et al., 2016). The biological mechanisms that impair chronic wound healing are still not fully understood, despite the advances made in wound care. Particularly, the comparative study on biological differences among various types of chronic wounds is limited. Since wound healing is a complex and dynamic process driven by multiple cell types and their intercellular communication, a high-resolution map of the cellular and molecular programs that either promote or inhibit proper healing process is necessary. Recent single-cell RNA sequencing (scRNA-seq) studies have begun to uncover the heterogeneity of stromal and immune cells in the chronic wound microenvironment. For example, recent studies have identified enrichment of a distinct fibroblast subset with high expression of inflammatory and ECM remodeling genes, inflammatory macrophage, and naïve T cell populations in healing DFU (Chen et al., 2022, Choi et al., 2025, Theocharidis et al., 2022). However, most studies have been focused on a single wound type or species, which made it difficult to obtain comprehensive view of the shared and distinct cellular programs across acute and diverse chronic wounds. Therefore, we have taken a cross-species and cross-pathological approach to address this gap in this study. We collected a large-scale single-cell data of wound tissues from both human (Choi et al., 2025, Davis et al., 2020, Januszyk et al., 2020, Justynski et al., 2023, Liu et al., 2025, Rustagi et al., 2022, Sandoval-Schaefer et al., 2023, Theocharidis et al., 2022) and mouse (Abbasi et al., 2020, Gay et al., 2020, Haensel et al., 2020, Ma et al., 2023, Pang et al., 2021, Phan et al., 2020, Wang et al., 2023), including healthy and diabetic skin, acute wounds, and chronic ulcers such as diabetic wounds and venous ulcers. This allowed us to capture the cellular and molecular landscapes of fibroblasts, macrophages, T cells, and other stromal and immune compartments. We systematically defined cell states, regulatory modules, and pathways associated with successful versus impaired wound healing by performing pseudotime analysis, transcription factor activity inference, and non-negative <u>matrix</u> factorization (NMF) analysis. To make these findings accessible for the wider research community, we developed WoundSCAtlas, an interactive web resource which offers visualization and comparison of cell states across species, wound types, and healing outcomes. Collectively, this study provides a comprehensive single-cell framework to better understand the cellular and regulatory basis of wound repair and offers a valuable resource for future mechanistic discovery and therapeutic development in chronic wounds.

## Results

### Establishment of a comprehensive and integrative cross-species single-cell atlas for wound biology

To address the heterogeneity and complexity of chronic wound pathogenesis and healing, we established the WoundSCAtlas, a large-scale single-cell transcriptomic resource that reveals conserved and species-specific cellular populations underlying wound healing process and failure. As summarized in the study workflow (**Fig. 1**), the atlas was created by assembling and integrating single-cell data from both patients and mouse models. The atlas comprises over 351,993 single cells from humans (Choi et al., 2025, Davis et al., 2020, Januszyk et al., 2020, Justynski et al., 2023, Liu et al., 2025, Rustagi et al., 2022, Sandoval-Schaefer et al., 2023, Theocharidis et al., 2022) including acute wound (AW, n=48,505), chronic ulcer (CU, n=48,473), diabetic foot ulcer (DFU, n=112,211), and chronic venous ulcer (CVU, n=19364), and 148,548 single cells from diverse mouse models (Abbasi et al., 2020, Gay et al., 2020, Haensel et al., 2020, Ma et al., 2023, Pang et al., 2021, Phan et al., 2020, Wang et al., 2023) including AW (n=69,863), chronic wound (CW, n=9,919), diabetic wound (DW, n=32,400), and infected wound (n=7,720). To facilitate community-wide exploration of the datasets and analytical results generated in this study, we developed WoundSCAtlas, a web-based interactive platform that integrates single-cell RNA-seq data from both human and mouse wound tissues. All processed data and analytical results were compiled into the WoundSCAtlas online tool (bhasinlab.bmi.emory.edu/WoundSCAtlas). This platform provides interactive visualization, pathway analysis, and gene correlation functionalities, enabling researchers to explore cell type specific gene expression patterns, predicted signaling pathways, and transcription factors (**Fig. 1**). The interface allows interactive visualization of UMAP embeddings, transcriptional module activity, transcription factor networks, and pathway enrichment, as well as query of gene-level expression patterns across cell types and wound from human and mouse models.

**Figure 1.**
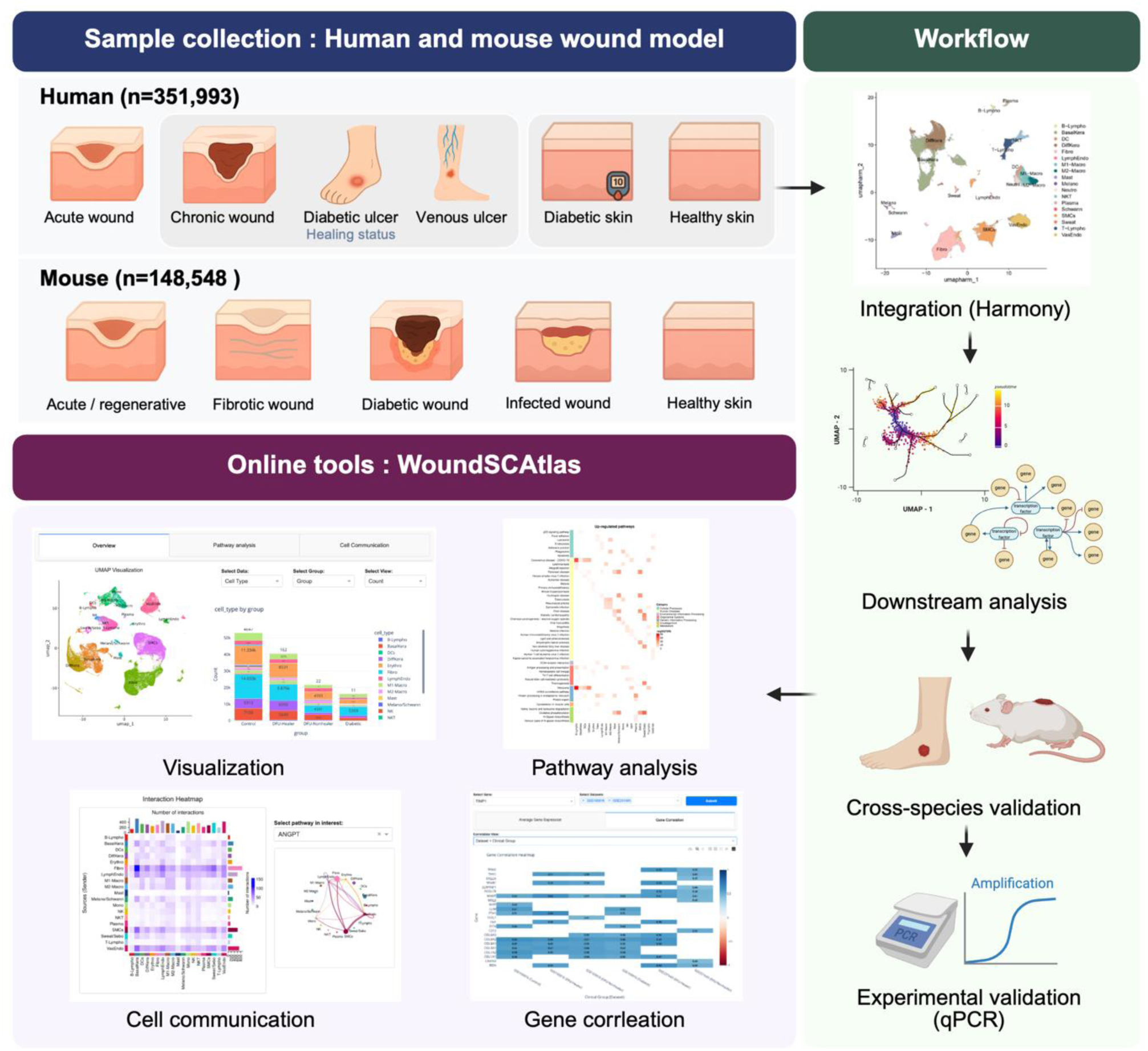
Overview of the wound single-cell atlas and analytical workflow. Human and mouse datasets were collected across diverse wound types, including acute, chronic, diabetic, and venous ulcers, alongside healthy and diabetic skin controls. Mouse models encompassing regenerative, fibrotic, and diabetic wounds were integrated to capture conserved and species-specific responses. After single-cell RNA-seq preprocessing, datasets were integrated, followed by downstream analyses including transcriptional modules discovery, pathway and transcription factor enrichment analysis, and intercellular communication analysis. Key findings were validated experimentally and across species to identify robust and conserved transcriptional programs. Finally, all datasets and analysis outputs were implemented into WoundSCAtlas (https://bhasinlab.bmi.emory.edu/WoundSCAtlas/), an interactive web resource that supports visualization, pathway analysis, cell-cell communication, and gene correlation exploration.

### Single-cell RNA-Seq analysis reveals conserved and divergent cellular remodeling in human and mouse wounds

To characterize the cellular heterogeneity and explore the regulatory changes across wound types and species, we compiled scRNA-seq data from both human and mouse dermal tissues (**Fig. 1, Table S1A, B**). Human samples included healthy skin, diabetic skin, AW, and chronic ulcer including DFU and CVU. We applied stringent quality control and obtained 351,993 cells from human and 148,548 cells from mouse (**Fig. S1A**). For each species, normalized and preprocessed gene expression matrix was used to perform dimensionality reduction by principal component analysis (PCA) and Unsupervised Uniform Manifold Approximation and Projection (UMAP), and clustering. The transcriptomically distinct cellular clusters were annotated based on the expression of established marker genes. We identified 18 canonical cell types in humans (**Fig. 2A, B),** out of which 16 also observed in the mouse models (**Fig. 2C, D**). These included cell types such as (‘Fibro’; human: *DCN+, PDGFRA+*; mouse: *Dcn+, Pdgfra+*), basal keratinocytes (‘BasalKera’; human: *KRT5+, KRT14+*; mouse: *Krt5+, Krt14+*), differentiated keratinocytes (‘DiffKera’; human: *KRT1*+, *KRT10*+; mouse: *Krt1*+, *Krt10*+), T lymphocytes (‘T-Lympho’; human: *CD3D+,* mouse: *Cd3d+*), M1-like macrophages (‘M1-Macro’; human: *IL1B+,* mouse: *Il1b+*), and M2-like macrophages (‘M2-Macro’; human: *CD163+*; mouse: *Mrc1+)* (**Fig. 2B, D**). All datasets were integrated using Harmony algorithm to adjust the batch effects due to technical variations. Cell abundance based comparative analysis depicted distinct cell type redistribution patterns associated with chronic wound pathogenesis in both species (**Fig. 2E, F**). These chronic signatures involved a significant loss of regenerative potential characterized by a decrease in keratinocyte populations(Senoo et al., 2007, Zhang et al., 2015). Specifically, human chronic ulcers (CU, DFU-NH, CVU) exhibited the reduced proportion of DiffKera, whereas mouse chronic models showed a prominent decrease in BasalKera. This keratinocyte loss aligns with the established role of keratinocytes that proper migration and proliferation of the cells are required for re-epithelialization and subsequent wound healing (Usui et al., 2008). Moreover, in human non-healing wounds, we observed persistent inflammation signs with a significant increase in both lymphoid lineage (T-Lympho) and myeloid lineage (neutrophils; Neutro) compartments as compared to healthy skin (**Fig. 2E**). Similar inflammatory signs were also observed in mouse chronic wound models (**Fig. 2F**). Furthermore, the proportion of Fibroblasts consistently showed a decreasing trend in human chronic ulcers (CU, DFU, CVU) compared to AW, and was notably lower in DFU-NH than in DFU-H. A similar pattern was observed in the chronic wounds of mouse models (**Fig. 2F**). This reduction in the key matrix-producing cell population, coupled with persistent inflammation, underscored the need for an in-depth characterization of the cell state heterogeneity of the dominant stromal and immune populations to identify the novel cellular subtypes driving pathophysiology impairing proper wound healing.

**Figure 2.**
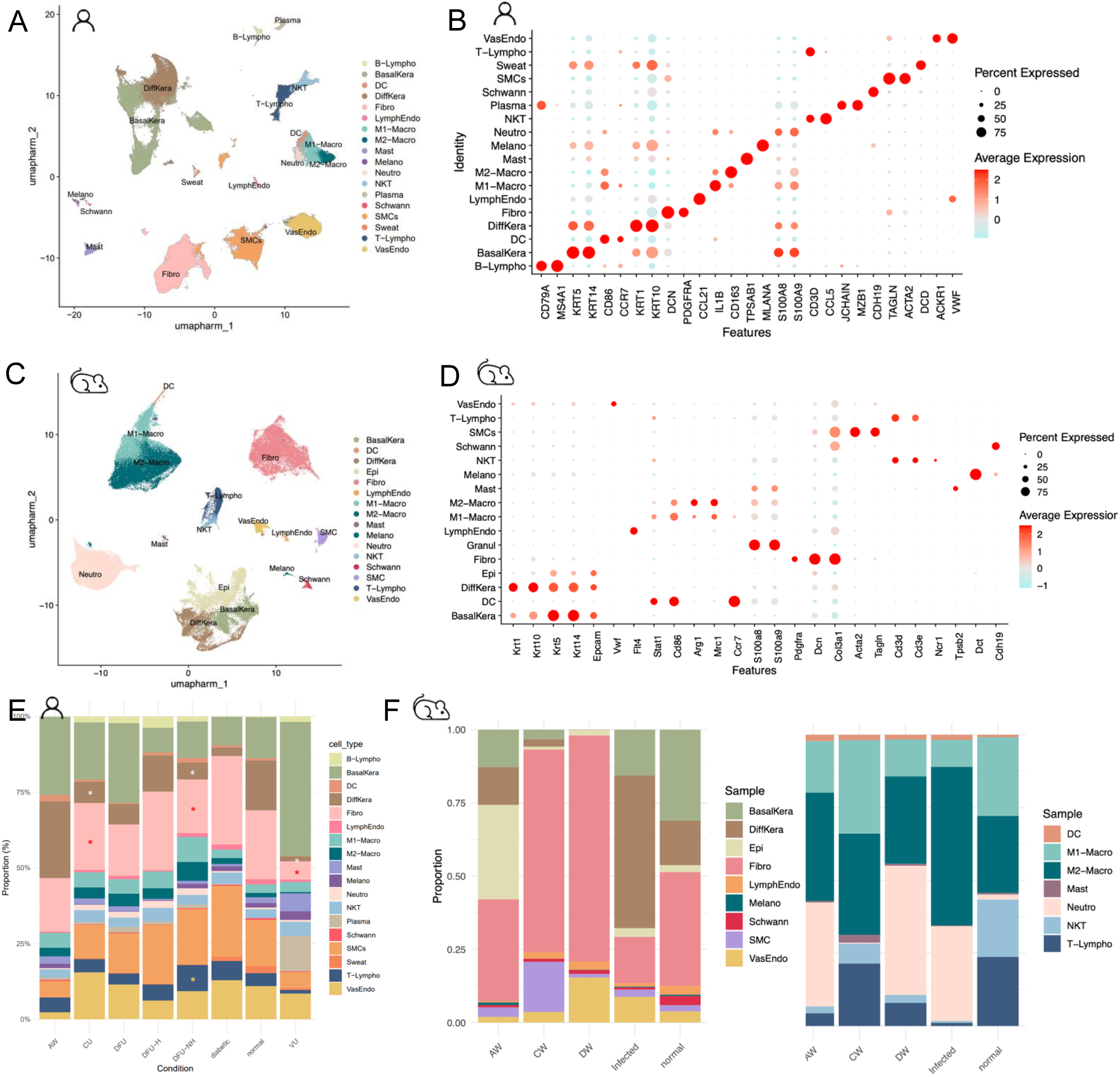
Overview of cellular composition and annotation of human and mouse wound single-cell atlases. **A.** UMAP embedding of all human dermal cells colored by annotated cell type, revealing diverse stromal, immune, and vascular populations within wound and skin samples. **B.** Dot plot showing expression of canonical marker genes used for cell type annotation in the human wound atlas **C.** UMAP representation of the mouse wound atlas, highlighting analogous cellular populations identified across murine wound and skin models. **D.** Dot plot depicting marker gene expression patterns used for annotation of the mouse atlas **E.** Proportional distribution of human cell types across wound conditions, demonstrating variation in cell types between acute, chronic, diabetic, and venous ulcer tissues. **F.** Cell type composition in the mouse datasets, with the summarizing non-immune (stromal, epithelial, and vascular) compartments (left) and the major immune cell populations (right), across healthy, acute, fibrotic, infected, and diabetic wound models.

### Fibroblast trajectory bifurcation reveals STAT3 as a potential regulator for healing inducive programs in chronic wounds

We conducted pseudotime trajectory analysis to investigate the continuous changes of cell states within Fibro populations (**Fig. 3A**). After sub-clustering, the cluster with progenitor-like transcriptional features, characterized by high *PDGFRA* and *CD44* expression combined with comparatively lower levels of *COL3A1* and *PDPN* was chosen as the trajectory’s root (**Fig. 3B, Fig. S2A-C**). Two distinct lineages were identified, bifurcating toward acute and chronic wound states. The acute lineage mainly consists of cells from AW with progression of trajectory correlated with the post-operative days (**Fig. 3C**, **Fig. S3A**). Conversely, there was a progressive shift from healthy, DFU-Non-healer to DFU-Healer and CVU cell fractions in chronic lineage (**Fig. 3C**). Differential expression analysis between two lineages exhibited that *IGFL3* and *IGFL2* were highly expressed in the acute lineage while inflammation related genes such as *CHI3L1*, *CXCL6*, and *CXCL13* were overexpressed in the chronic wound lineages (**Fig. 3D, Table S2**). We further investigated genes that are specific to each lineage through grouping them by their expression patterns along pseudotime and performed pathway enrichment analysis. In the acute lineage, the regulation of IGF transport and uptake by IGFBPs was upregulated in the early stage. This process preceded Wnt signaling, collagen formation, epidermal cell development pathways, and followed by extracellular matrix and structure organization (**Fig. S3B**). However, genes upregulated in the early stage of the chronic lineage were significantly associated with negative regulation of TCF-dependent signaling by WNT ligand antagonists. Subsequently upregulated genes were involved in NF-kB signaling pathway, signaling by NTRKs and interleukins (**Fig 3E**), suggesting upregulation of inflammatory responses(Simmonds and Foxwell, 2008). These inflammation related pathways were prolonged in the later phase with JAK-STAT signaling pathway as well as chemokine signaling pathway that has been reported previously (Hu et al., 2021, Xue et al., 2023). To dissect the underlying regulatory landscape, we performed transcription factor (TF) activity inference within the two lineages. This identified STAT3, RELA, and CEBPB as the top TFs showing higher activity in the chronic lineage as compared to the acute lineage (**Fig. S4A**). As CEBPB and NF-kB family members are known players of inflammatory and stress-related programs (Cappello et al., 2009, Ren et al., 2023, Stein et al., 1993), they are likely to extend the inflammation phase in chronic wounds. Notably, while these TFs were significantly enriched within the chronic lineage, STAT3 activity uniquely distinguished the healing outcomes of chronic wounds. Specifically, the activity score of STAT3 regulon was higher in DFU-healer cells as compared to non-healers. Additional analysis using STAT3 gene set enrichment also recapitulated this finding with significantly higher enrichment scores in the healing chronic wounds (**Fig. 3F, G**). Particularly, STAT3 regulated *IL6*, *CHI3L1*, and *MMP3*, which constitute the signature gene sets of healing enrichment fibroblasts (HE-Fibro) defined in our previous studies (**Fig. 3H, 4SB-D**) (Choi et al., 2025, Theocharidis et al., 2022). This supports STAT3 as a potential key player of chronic wound reparative microenvironments, involved in inflammatory responses as well as ECM remodeling. Furthermore, these findings align with recent studies demonstrating that STAT3 activation promotes fibroblast proliferation, migration, and chronic wound healing via mitogen-activated protein kinase (MAPK) pathway, mediated by *CHI3L1* (Zhang et al., 2024, Zhao et al., 2020). We have also validated our findings in Fibro from murine wound models. Notably, the bifurcation of trajectories into acute and chronic lineage was also observed in mouse Fibro (**Fig. S5A, B**). Stat3, Rela, and Stat1 were among the top TFs exhibited high activity scores in chronic wound lineage, consistent with their human counterparts (**Fig. S5C**). Further in this lineage, we observed high expression of *Hif1a* that is regulated by STAT3 by binding its promoter, *Il6*, a cytokine that activates the STAT3 signaling pathway through JAK kinases, and *Mmp3*, a target gene of STAT3 involved in ECM remodeling (**Fig. S5D, E**). In summary, the fibroblast trajectory analysis showed fibroblast bifurcation toward acute and chronic wound states in both human and mouse wounds, and the activation of STAT3 could be a potential switch to healing enriched fibroblasts in chronic wounds.

**Figure 3.**
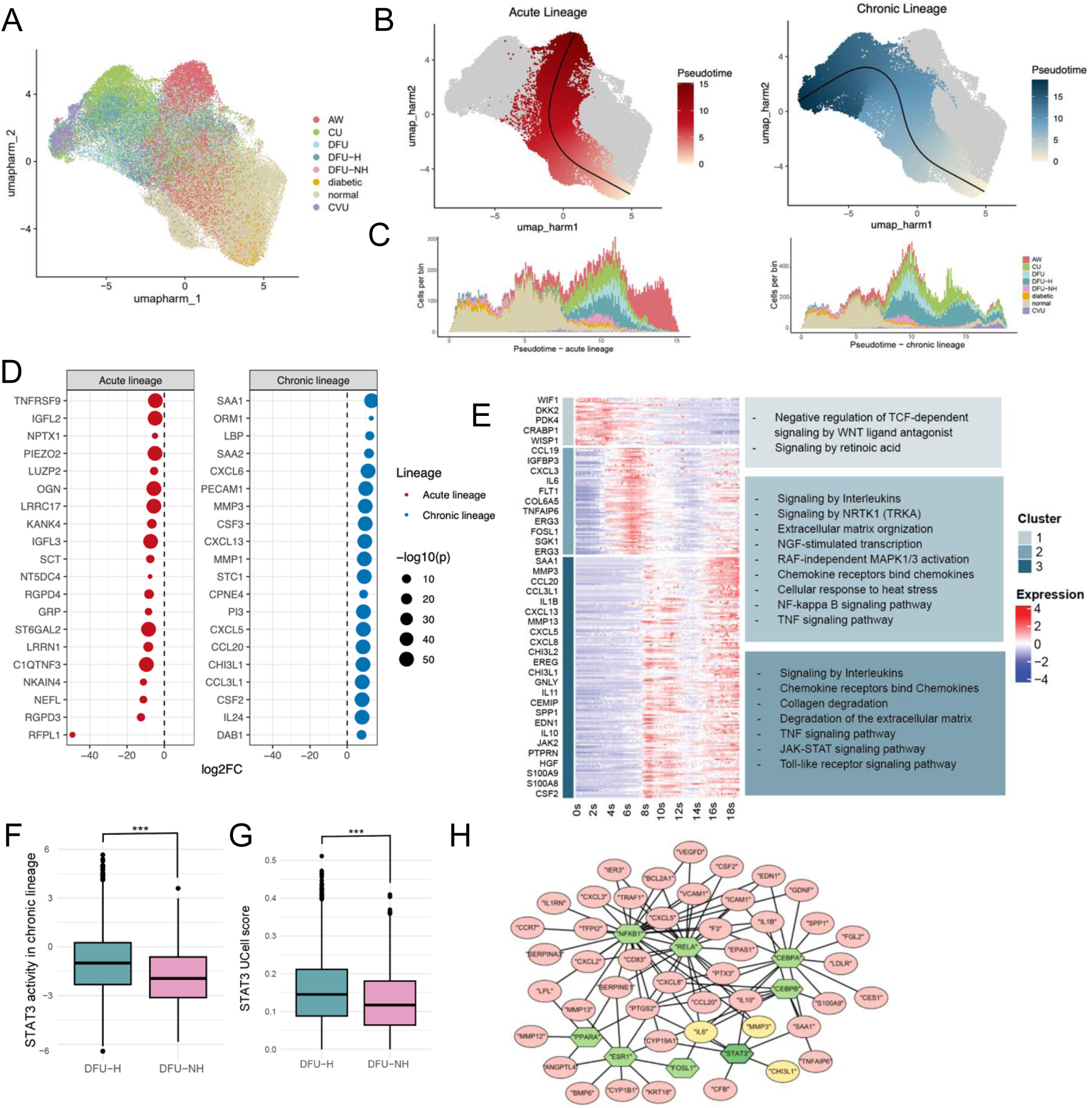
Fibroblast bifurcation reveals divergent transcriptional programs in acute and chronic wounds. **A.** UMAP visualization of fibroblasts across wound types. **B.** Pseudotime trajectories inferred by slingshot identify two major fibroblast lineages: an acute lineage associated with regenerative healing and a chronic lineage. **C.** Sample composition plotted along pseudotime demonstrates that acute lineage cells are enriched in acute and healing wounds, while chronic lineage progressively accumulates fibroblasts from chronic wound contexts, including DFU and VLU. **D.** Representative DEGs defining each trajectory. The acute lineage is enriched for genes associated with connective tissue organization and contraction, whereas the chronic lineage shows induction of stress- and inflammation-related genes (*MMP3, CXCL9, CHI3L1, IL6*). **E.** Heatmap showing the expression of the chronic lineage specific genes along the pueudotime and enriched pathways by groups **F.** STAT3 transcription factor activity scores showing higher activity in fibroblasts from healed DFU compared with non-healed DFU (***, p-value < .001). **G.** Ucell score of STAT3 target genes across pseudotime demonstrates preferential enrichment in the chronic fibroblast lineage (***, p-value < .001). **H.** Gene regulatory network highlighting top transcription factors and their target genes in chronic lineage.

### Chronic wounds contain exhausted regulatory T cells, whereas microenvironments of healing wounds are enriched for quiescent memory T cells

To investigate T-Lympho and natural killer T cells (NKT) heterogeneity across wound types, we subclustered these populations and applied NMF to identify transcriptional modules (**Fig. 4A, B**). This analysis identified distinct programs that mapped onto different wound types and outcomes. We identified two modules that were enriched in T-Lympho (T1, T2) and one (T3) in NKT. Specifically, the T1 module that is associated with DFU-H was a quiescent, memory-like program marked by upregulation of *FOXP1*, *FOXO1*, *TCF7*, related to the maintenance of T cell potential and long-term functionality (Kerdiles et al., 2009) (**Fig. 4C-E**). In contrast, the T2 module representing DFU-NH was characterized by high expression of inhibitory receptors such as *CTLA4*, *TIGIT*, together with regulatory T cell markers *FOXP3*, *IL2RA*, and *IFNG (Tang and Bluestone, 2008)* (**Fig. 4C-E**). Pathway enrichment analysis was performed based on genes that contributed to each module. Quiescent memory-like T cells showed enrichment of pathways such as regulation of localization of FOXO transcription factor and FLT3 signaling, which is crucial for survival and stress resistance (Kerdiles et al., 2009, Scheijen et al., 2004), while exhausted T cells and Tregs showed upregulation of TNFR2 non-canonical NF-kB pathway and interferon signaling pathways (**Fig. 4F**). These findings were further supported by inferred transcription factor activity and gene regulatory networks (GRNs). Exhausted T/Tregs exhibited increased SATB1 and MSC activity, which is linked to the T cell exhaustion markers, whereas quiescent T cells exhibited elevated FOXP1 and FOXO1 activity (**Fig. 4G, H**). T3 module with strong representation of CVU depicted increased expression of exhaustion associated markers including *LAG3* and *TIGIT*, suggesting that NKT may also undergo dysfunction in the context of chronic ulceration (**Fig. 4E**). In addition, T3 module depicted enrichment of PD-1 signaling pathway and translocation of ZAP-70 to immunological synapse (**Fig. 4F**). Finally, cross-species analysis demonstrated a couple of modules that were specific to a certain experiment model (**Fig. S6A**). MT1 module which was commonly associated with all diabetic wounds showed high *Il2rb* and *Nfkb1* (**Fig. S6B**). Particularly in murine type 2 diabetes wounds, we observed that exhausted T cells and Tregs module (MT3) was enriched with increased Foxp3, Tox, and other inhibitory receptors, paralleling the human T1 program. MT2 module was highly correlated with *Lgals3*, *Ctla2a*, and effector gene *Nkg7*, indicating enhanced cytotoxic potential in infected wounds. In conclusion, these conserved results across species indicate that T cells from healed chronic wounds maintain a quiescent, regenerative potential, while non-healing chronic wounds accumulate exhausted T/Tregs and putatively dysfunctional NKT cells, which could limit immune competence and potentially contribute to poor outcomes.

**Figure 4.**
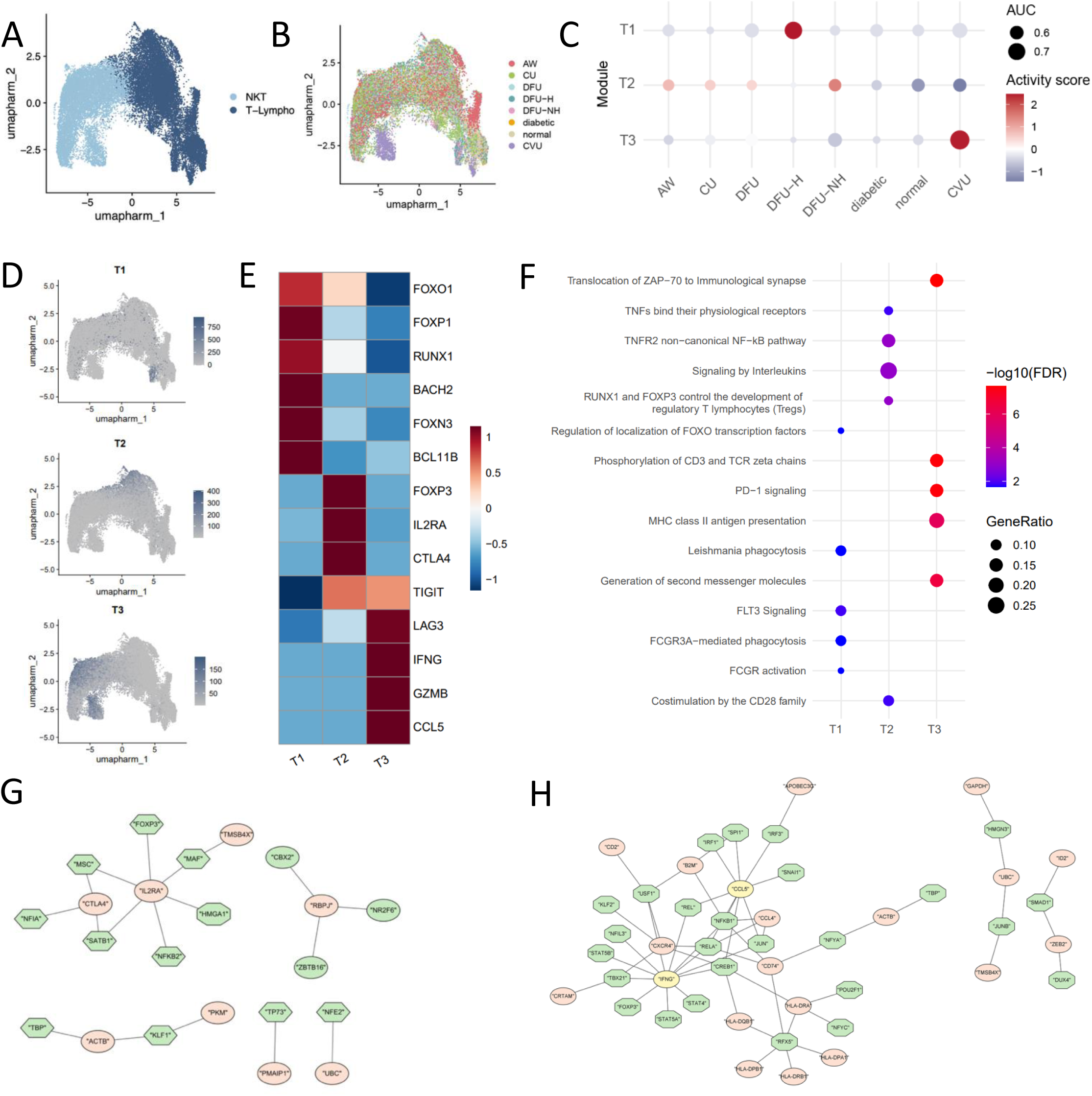
Divergent T cell states reveal a shift from quiescence to exhaustion from healing DFUs to non-healing DFUs. **A.** UMAP visualization of the T/NKT compartment showing distinct clusters corresponding to conventional T lymphocytes and NKT cells. **B.** Cells are colored by wound condition, revealing the distribution of T and NKT subsets across acute, diabetic, and venous ulcer wounds. **C.** Module activity scores across wound conditions showing enrichment of quiescent T cell programs (W6) in healed wounds, and exhaustion-related programs (W13, W4) in non-healed diabetic foot ulcers (DFU–NH) and venous ulcers (VLU). **D.** UMAP plots showing spatial distribution of T cells and NK cells transcriptional modules identified by non-negative matrix factorization. **E.** Heatmap showing representative genes defining each transcriptional module. **F.** Pathway enrichment analysis showing top pathways across selected modules. Gene regulatory network (GRN) reconstruction highlighting transcriptional regulators and their target genes in **G.** quiescence T cells from healing DFU and **H.** exhausted T regs from non-healing DFUs.

### Transcriptional divergence of macrophages uncovers pro-fibrotic SPP1 signaling and foamy phenotypes in chronic wounds

To decipher role of macrophages in wound outcomes, we performed focused analysis on myeloid compartments (M1-Macro, M2-Macro, Neutro, and DCs) and identified multiple transcriptional modules using NMF representing various functional states (**Fig. 5A, B).** Within the M1-Macro compartment, two distinct modules were identified (MW1 and MW2) **(Fig. 5C)**. AW specific module MW1 was characterized by overexpression of pro-inflammatory genes *TIMP1*, *IL1RB*, *CXCL8*, *CCR7* as well as antigen-presenting HLA class || molecules (**Fig. 5D, S7B**). In contrast, DFU-NH specific module MW2 was correlated with *SPP1*, *APOC1*, and *FTL* (**Fig. 5D, S7C**). Interestingly, recent studies defined SPP1+ macrophages as a profibrotic macrophages subtype that drives fibrosis by interacting with myofibroblasts through signals such as CXCL4 and IL-17A (Hoeft et al., 2023, Jiang et al., 2024). Following pathway enrichment analysis revealed that MW1 was highly enriched for T cell activation related pathways, including translocation of ZAP-70 to immunology synapse (Fernandez-Aguilar et al., 2023). It was also correlated with phosphorylation of CD3 and TCR zeta chains, which is known to trigger proinflammatory responses such as NF-kB signaling pathway (Zinatizadeh et al., 2021), while MW2 showed significant enrichment for inflammatory and metabolic dysfunction related pathways, for example, neutrophil degranulation and iron uptake and transport. Additionally, MW3 and MW4 modules were enriched with M2 Macrophages with enrichment of cells from CVU and AW respectively. MW3 module showed high expression of *SPP1* and lipid metabolism-related genes such as *APOE*, *APOC1*, and *LIPA* (**Fig. 5E, S7D**) (Tudorache et al., 2017). Further pathways analysis showed enrichment gene set for binding and uptake of ligands by scavenger receptors, suggesting association with foam cell formation (**Fig. 5G**). According to recent studies, foamy macrophage, which are characterized by high expression of lipid-processing genes, has reduced migration ability and a higher chance of apoptosis (Poznyak et al., 2021). One study suggested that foam-like macrophages delay wound healing by preventing fibroblasts from migration, proliferation, and chemokine secretion through interfering TLR/NF-kB signaling (Wang et al., 2022). In contrast, MW4 which was enriched in acute wounds, carried significant enrichment of *CXCL* chemokines which is crucial for immune cell recruitment and heat shock proteins (**Fig. 5E, G**). Further, cell-cell communication analysis revealed that the *SPP1*+ M1-Macro of DFU-NH and the *SPP1*+ M2-Macro of CVU were dominant sources of SPP1 signaling pathway (**Fig. 5H**). Cross-species validation also demonstrated that the foamy macrophage phenotype was conserved in type 2 diabetes wound models, with high expression of *Abca1, Fabp5, and Trem2*, recapitulating our observations in human MW3 module (**Fig. S8A**). Taken together, these results established that macrophages states vary within the wound environment. Specifically, we observed healer associated *TIMP1*+ M1-Macro in contrast to non-healer associated *SPP1*+ M1-Macro, and foam-like M2-Macro enriched in CVU and DFU.

**Figure 5.**
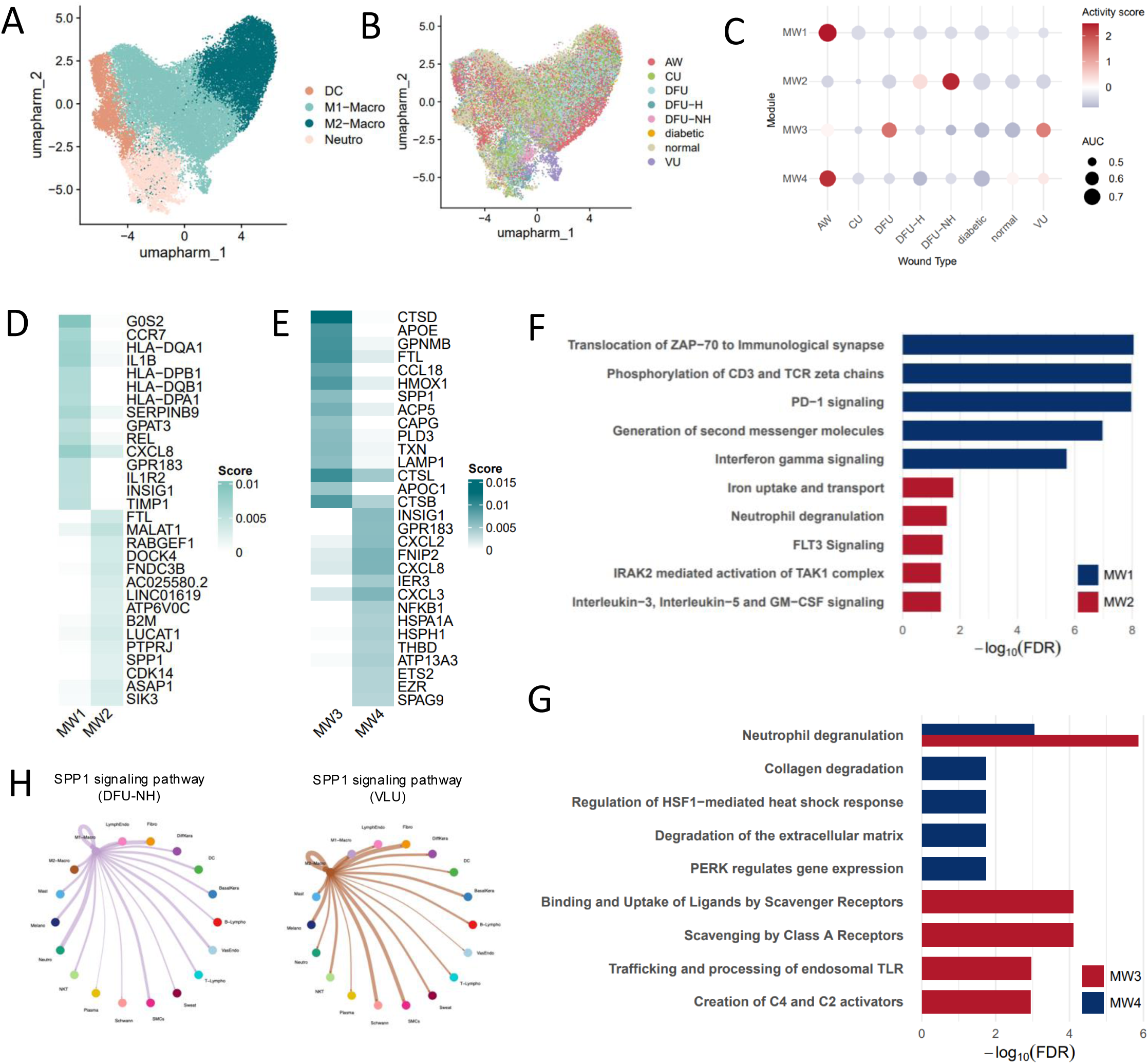
Distinct myeloid programs delineate healing-associated TIMP1⁺ M1 macrophages and SPP1⁺ macrophages in chronic wounds. UMAP visualization of myeloid cells from all wound types, highlighting **A.** dominant cell types including M1-Macro, M2-Macro, and DC **B.** wound types. **C.** Module activity scores across wound types reveal selective activation of TIMP1⁺ M1 module in healed DFU and SPP1⁺ macrophage module in non-healed DFU and VLU. Heatmaps showing representative top-contributing genes for each module in **D**. M1-Macro and **E.** M2-Macro. Pathway enrichment analysis results based on highly correlated genes to transcriptional modules of **F**. M1-Macro and **G.** M2-Macro. **H.** Ligand–receptor interaction analysis demonstrates activation of SPP1-mediated signaling in DFU-NH and VLU, implicating osteopontin pathways in chronic wound persistence.

### Machine learning models identified a biomarker for chronic wound outcome and its clinical validation via RT-PCR

To identify a molecular gene signature that distinguishes DFU-H and DFU-NH, we aggregated expression profiles of scRNA-seq data into pseudo-bulk compartments. These represent cell types of Fibro, SMCs, M1-Macro, and additional stromal and immune lineages (**Fig. S9A**). We trained multiple machine learning classifiers on psuedobulked matrices to classify healing and non-healing outcomes. Using feature importance ranking, we identified a combination of three genes that were consistently ranked highly and acquired plateaued classification accuracy (**Fig. S9B**). These include *CHI3L1*, *TIMP1*, and *SPP1,* and they achieved ROC-AUC of 0·805 in 5-fold cross validation (**Fig. 6A**). It was shown that *TIMP1* and *CHI3L1* were strongly associated with HE-Fibro, SMCs, and M1-Macro with SVM coefficients of 1·1046 and 0·7219, respectively, whereas *SPP1* was enriched in M1-Macro in non-healing contexts having a negative coefficient of 0·5520. To evaluate the ability to translate signature genes’ measurements into clinical and experimental environments, we measured their expressions by RT-qPCR on wound dressings collected from patients with DFUs (**Fig. 6B**). Wound outcomes were determined by a complete wound closure after 12 weeks from the first visit. Subsequently, our SVM model classified RT-qPCR measured expression of the genes accurately discriminated against healed and not healed DFU samples with the accuracy of 85·71%, recall of 83·33%, specificity of 86·67%, and ROC-AUC of 0·844 (**Fig. 6C**). Further validating the discriminatory power of this gene signature supports its potential use as a clinically translatable biomarker for predicting chronic wound outcomes.

**Figure 6.**
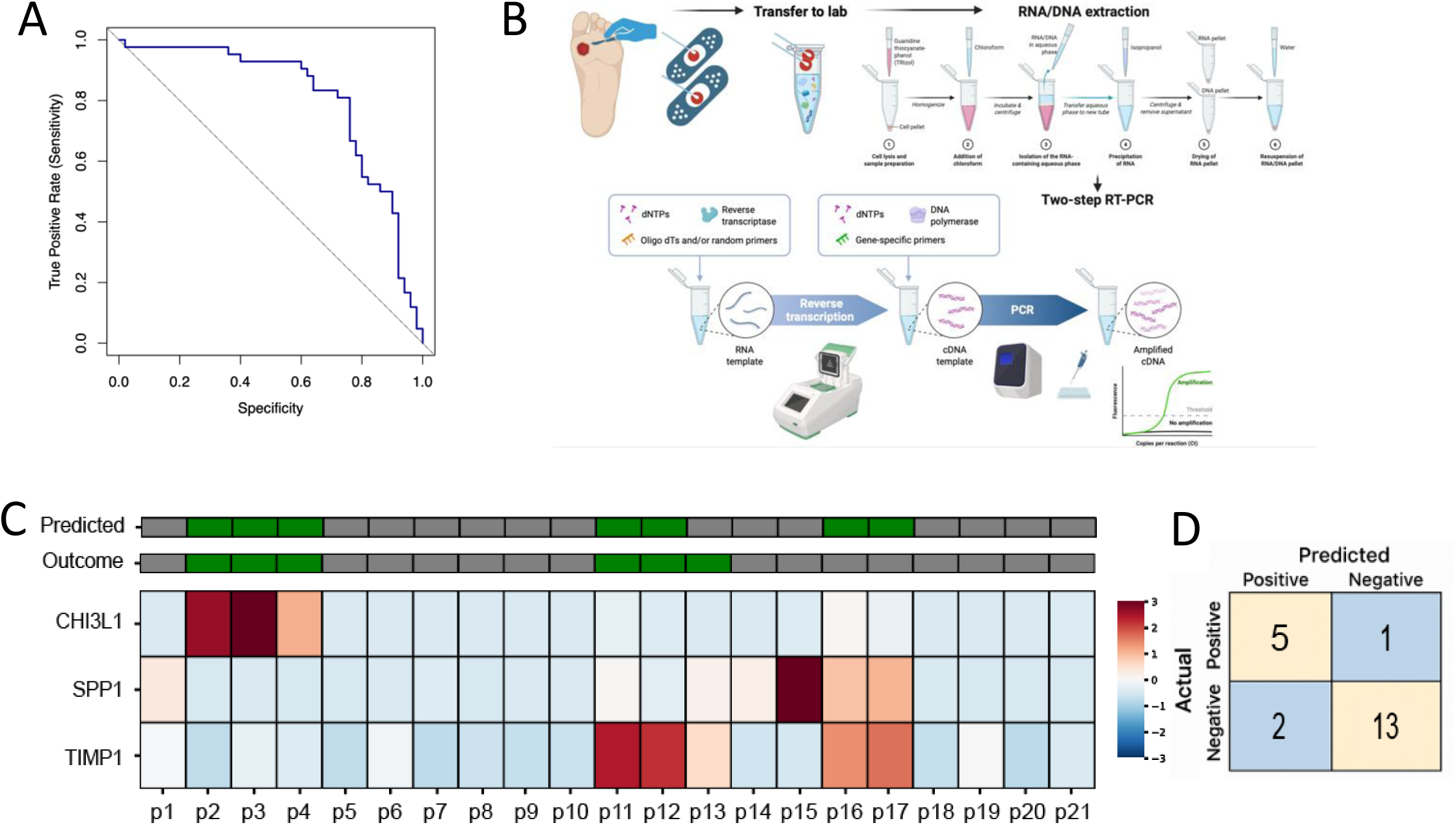
Validation of healing prediction gene set in RT-qPCR. **A.** ROC curve demonstrating performance of healing prediction genes (*CHI3L1, TIMP1, SPP1*) on a test set of pseudobulked expression matrix. **B.** Workflow for processing diabetic wound bandages including sample collection, RNA extraction, and RT-qPCR. **C.** Heatmap showing the scaled expression of RT-qPCR values for three key biomarkers across 21 individual wound bandage samples. The top annotation bars display the predicted healing outcomes by SVM model and the actual clinical outcomes (green: healer, grey: non-healer). **D.** Confusion matrix summarizing the performance of SVM model based on LOOCV.

## Discussion

In this study, we generated a comprehensive single-cell transcriptomic atlas, encompassing multiple wound types and healing outcomes from both human and mouse. By analyzing 351,993 cells of human and 148,548 cells of mouse, we delineated conserved molecular and cellular programs that lead chronic wound healing and prolonged inflammation. We identified two distinct fibroblast lineages corresponding to acute and chronic wound states by trajectory analysis. Importantly, among chronic lineage specific TFs, we observed that STAT3 activation might functions as a key regulator of fibroblasts to drive chronic wounds healing. Elevated STAT3 activity was associated with the overexpression of healing-enriched fibroblasts signature genes, specifically *CHI3L1*, *IL6*, and *MMP3*. These conserved fibroblast trajectories and transcriptional regulators in murine diabetic and infected wound models underscored our findings in human fibroblast compartment. Within the immune compartment, we identified functional discrepancies within both T cells and macrophages across wounds that are destined to be healed or not. T cells in healing chronic wounds exhibited a quiescent, memory-like phenotype marked by *FOXO1* and *FOXP1*. Whereas non-healing chronic wounds accumulated exhausted T-Lympho expressing *CTLA4* and *LAG3* and T regulatory cells, which were associated with upregulation of PD-1 signaling, implicating chronic antigenic stimulation in impaired immune responsiveness. These findings suggested that preservation of immune quiescence, rather than sustained immune activation, may be permissive for effective tissue regeneration. In myeloid compartment, we identified a *TIMP1*+ M1-like macrophages enriched in healing wounds, expressing immune-stimulatory and matrix remodeling factors, in contrast to *SPP1*+ macrophages in non-healing ulcers, exhibiting foam-like features such as *APOE*, *APOC1*, and lipid metabolism pathways. Notably, *TIMP1*, *CHI3L1*, and *SPP1* reflected distinct fibroblast and macrophage states that were correlated with healing status of chronic wounds and identified by machine learning models to discriminate healing outcomes. RT-qPCR measured gene expressions from wound dressings accurately classified healing and non-healing DFU samples using the SVM classifier. This demonstrated a clinical feasibility of healing gene signature beyond single-cell discovery. Finally, the WoundSCAtlas, which is directed to enable exploration of transcriptional modules, pathway, and intercellular communication across multiple datasets, bridges experimental and clinical research, and may guide target discovery for potential therapeutic targets. Future studies utilizing spatial transcriptomic profiling are warranted to see if there are a spatial proximity and organization of immune cells and healing associated fibroblasts. In addition, prospective validation of the healing gene signature in larger and independent patient cohorts is necessary for better robustness. In conclusion, this work delineated a cellular and molecular landscapes of wound healing and chronicity, exhibiting cross-species conservation on immune exhaustion, macrophage plasticity, and fibroblast reprogramming.

## Methods

### Sample collection and preprocessing

Single-cell RNA-seq datasets from both human and mouse dermal tissues were curated from the Gene Expression Omnibus (GEO). Dataset encompasses diverse wound conditions, including acute wounds, chronic wounds, diabetic foot ulcers, and venous ulcers and their healing status. It contains murine excisional wound healing models. Accession numbers, sample meta data, and detailed information are listed in **Table S1.** Cells were filtered to remove low-quality cells and doublets across all datasets with the thresholds of <200 detected genes, <15% mitochondrial reads, and <25,000 UMIs. Potential doublets were identified with DoubletFinder(McGinnis et al., 2019) and removed. Cells expressing <200 genes and genes expressed in <3 cells were excluded. Each dataset was normalized, log-transformed, scaled by Seurat (v5)(Hao et al., 2024).

### Data integration and batch correction

Within each species, all datasets were merged by R merge function. We used Harmony (v1.2)(Korsunsky et al., 2019) to correct for batch effects derived from different studies, while preserving biological variation. 3,000 highly variable genes (HVGs) identified by FindVariableFeatures were used for downstream analysis. Dimensionality reduction was performed using PCA followed by UMAP. Clusters were identified using FindNeighbors, the shared nearest neighbor (SNN) graph-based modularity algorithm.

### Non-negative matrix factorization (NMF) and module discovery

For each major lineage, gene programs were identified using Bayesian Non-negative Matrix Factorization (NMF) implemented in SignatureAnalyzer_GPU package(Taylor-Weiner et al., 2019). Matrices were prepared specifically for NMF. For each cell compartment, raw count matrices were preprocessed using SCTrasnform to stabilize variance, followed by RMPK normalization. The algorithm decomposes the input matrix X into two non-negative components, W and H, representing gene weights and module activity weight, respectively. We adopted an L1 prior for W to promote gene sparsity within modules, while using an L2 prior to H to stabilize per-cell activity estimates. We ran optimization for up to 100,000 iterations or until convergence. The optimal number of factors (k) was selected automatically by the hierarchical automatic relevance determination (ARD) criterion. Genes with the top 50 weights per module were used to perform Reactome pathway enrichment using clusterProfiler and ReactomePA packages (adjusted *p*<·05)(Milacic et al., 2024, Wu et al., 2021). H matrix was mapped back to the Seurat object and normalized within wound type, enabling comparison of module activity.

### Trajectory inference

Pseudotime trajectories were perfomed using Slingshot (v2.12.0) on the UMAP embedding. Lineage roots were set to progenitor-like cluster (PDGFRA⁺, COL3A1⁻, CD44⁺). Gene expression dynamics along each lineage were modeled using tradeSeq (Van den Berge et al., 2020) (v1.18.0) which uses generalized additive models (GAMs) to identify genes showing significant changes over pseudotime and to compare genes having different patterns across lineages. Genes with adjusted *p*<·05 were grouped via hierarchical clustering and pathway enrichment was performed on grouped gene sets using clusterProfiler with Reactome and KEGG databases.

### Cell-cell communication analysis

Intercellular signaling was inferred per species and per wound type using CellChat(Jin et al., 2021) (v2.1.2). Ligand-receptor databases (CellChatDB-human/mouse) was used to calculate communication probabilities between annotated cell types. Communication strength was estimated using the computeCommunProb, and pathway level aggregation was done by computeCommunProbPathway. Differential communication analysis was performed across wound types as well as healing outcomes to identify wound specific ligand–receptor pathways.

### Healing enrichment gene set selection

To identify a gene signature that can predict chronic wound healing outcome, we implemented a multi-step feature selection framework. Candidate genes were initially drawn from the set of HVGs. We applied LASSO logistic regression to the healing dataset to shrink coefficient and select genes most associated with healing (positive) vs. non-healing (negative) cell identify. Positive cells were defined as fibroblasts, SMCs, and M1-macrophages from healed or healing samples (n=21,425) whereas negative cells included all remaining cell types (n=42,617). The selected features were used to train three independent classifiers: support vector machine (SVM), random forest (RF), and XGBoost on 80% of the data with the remaining 20% reserved as test set. Models were evaluated using ROC-AUC. Feature importance was interrogated using Shapely Additive exPlanations (SHAP) values derived from the trained classifiers. Genes were ranked by their absolute SHAP contribution to the prediction. To construct the final healing enrichment gene set, we iteratively added genes in descending SHAP order and assessed classification performance on training set. The final cutoff corresponded to the minimal number of genes that achieved a plateau in performance was evaluated on test set.

### Extraction of RNA from Wound Bandage

After receiving the wound bandages on ice, the blood and tissue residues were cut and transferred into RNA lysis buffer supplemented with 2-mercaptoethanol (RLT+2BME). Samples were vortexed briefly to ensure proper mixing, then incubated at room temperature for 10 minutes. Following incubation, the samples were centrifuged at 1000g for 10 minutes, and the supernatant was collected for RNA extraction using the Quick DNA/RNA Miniprep Plus Kit (Zymo Research, Cat. No. D70033) according to the manufacturer’s instructions. RNA concentration and purity were determined by using NanoDrop. First-strand cDNA was synthesized using the 4X ABScroipt Neo RT kit (ABclonal, Cat. No. RM21486) following supplier’s protocol.

### Quantitative Real-time PCR (qPCR)

RT-qPCR was performed to quantitatively measure gene expression levels in the samples with high sensitivity and accuracy. Quantitative qPCR was performed using 2X Universal SYBR green qPCR master mix (ABclonal, cat. No. RM21203). Amplification reactions were carried out in the Bio-Rad CFX Opus 96 real-time PCR system. Each reaction was prepared in a final volume consistent with the manufacturer’s recommendations and contained SYBR green master mix, gene-specific primers, and diluted cDNA. The housekeeping gene ACTIN was used as an internal control for normalization. PCR primers were designed using the NCBI Primer-Blast tool to specifically amplify the target sequences. The designed primers were checked for specificity and optimal melting temperature. After finalizing the sequences, primers were synthesized and ordered from Integrated DNA Technologies (IDT, USA). The primer sequence used in this study was as follows:

ACTIN: Forward: GAATCAATGCAAGTTCGGTTCC Reverse: TCATCTCCGCTATTAGCTCCG

CHI3L1: Forward: AAGCAACGATCACATCGACAC, Reverse: TCAGGTTGGGGTTCCTGTTCT

TIMP1: Forward: CTTCTGCAATTCCGACCTCGT, Reverse: ACGCTGGTATAAGGTGGTCTG

SPP1: Forward: GAAGGGAAAGTTGTCAGCGAA, Reverse: CCCCTGGTCTGTTGACTATTTG

RT-qPCR cycling (40 cycles) conditions were set according to the SYBR green mix manufacturer’s instructions. Relative gene expression levels were calculated using the 2^-ΔΔCT^ method (Livak and Schmittgen, 2001), with ACTIN as the reference gene.

### Prediction of wound healing outcome using qPCR data

Fold change of signature genes was scaled by standard normalization and used as covariates of the following analysis. To deal with class imbalance issue, we trained a linear SVM with class_weight=’balanced’. We employed a leave-one-out cross-validation (LOOCV) strategy due to limited available patient samples, and all metrics (accuracy, sensitivity, specificity, and ROC-AUC) were averaged across folds. Linear SVM coefficients were extracted to assess the relative contribution and directionality of the signature genes to outcome prediction.

### WoundSCAtlas web resource

We developed WoundSCAtlas (https://bhasinlab.bmi.emory.edu/WoundSCAtlas), an interactive web resource containing single-cell RNA-seq datasets of wound tissues. All datasets underwent a standardized preprocessing pipeline (data quality, log normalization, clustering, and cell type annotation). For datasets with no accessible cell type labels, reference-based cell type annotation was performed. Users can explore datasets by wound types or species, visualize UMAP embeddings, cell-type composition, cell-cell communication, pathway, and transcription factor enrichment analysis results.

### Patient and ethics statement

This study involved previously published and publicly available single-cell transcriptomic datasets that had received approval from respective Institutional Review Boards (IRB).

## Funding

This work was supported by Emory University Startup funds to MKB.

## Contributors

DC and MKB designed and coordinated the study. Clinical coordination and surgical procurement were performed by JM. Laboratory work was done by MB and AA. Bioinformatic analysis and visualization were performed by DC. DC drafted the manuscript and figures. Manuscript was reviewed and revised by MB, SB, MKB. All authors read and approved the final manuscript.

## Notes

### Competing Interest Statement

The authors have declared no competing interest.

